# CARTnet: Template-guided B-cell Epitope Prediction with Contact-aware Attention

**DOI:** 10.64898/2026.11.03.764112

**Authors:** Raj S. Roy, A N M Nafiz Abeer, Byung-Jun Yoon, Xiaoning Qian

## Abstract

B-cell epitope prediction remains challenging due to the limited availability of experimentally characterized data and the incomplete understanding of antibody–antigen recognition. Here, we present CARTnet, a novel contact-aware deep learning framework that integrates multimodal features and template-derived epitope information. CARTnet demonstrated superior performance when compared to the existing state-of-the-art methods across most evaluation metrics on both the internal and external test datasets. In particular, CARTnet achieved an AUPRC of 0.243 on the internal test set, compared with 0.220 for DiscoTope-3, 0.140 for BepiPred-2, and 0.198 for BepiPred-3. This improvement was also observed on the external test set, where CARTnet achieved an AUPRC of 0.267, compared with 0.192, 0.111, and 0.215 for DiscoTope-3, BepiPred-2, and BepiPred-3, respectively, supporting its generalizability. CARTnet also supports template-guided prediction, in which limited prior epitope information can improve the identification of additional epitope residues; a single template residue increased the AUPRC of the template-based variant from 0.225 to 0.403 on the internal test set. Furthermore, we demonstrated that epitope information from closely related viruses can be transferred across the genus Alphavirus to improve epitope prediction accuracy. These results establish CARTnet as an accurate framework for both *de novo* and template-guided B-cell epitope prediction.

**Data and Code Availability:** The source code and implementation of CARTnet are publicly available on GitHub at https://github.com/RajShekhorRoy/CARTnet.

## 1 Introduction

B-cell mediated humoral immunity plays a central role in host defense against emerging pathogens [1, 2] through the production of antigen-specific antibodies. Antibodies recognize distinct regions of antigens known as B-cell epitopes, and the identification of these epitopes is important for elucidating antigen–antibody interactions and facilitating vaccines and other therapeutic development. Consequently, accurate identification of B-cell epitopes has become a critical area of immunological and bioinformatics research.

Traditionally, B-cell epitopes were experimentally identified using techniques such as peptide arrays, ELISA, mutagenesis, and X-ray crystallography [3–5]. Although these approaches provide valuable experimental evidence, they can be time consuming and resource intensive [6, 7]. Computational B-cell epitope prediction therefore provides an efficient alternative approach for prioritizing potential antibody-binding regions and reducing the experimental search space for subsequent validation.

To address this limitation, many *in silico* tools have been developed. Early computational methods for predicting linear B-cell epitopes have identified highly hydrophilic regions using sliding-window approaches, based on the assumption that antibody-binding residues are likely to be surface-exposed and located in hydrophilic regions of proteins [8]. Other related approaches including ElliPro [9] exploit the geometric protrusion of residues from an ellipsoid fitted around the protein to predict epitopes. Also, empirical epitope information has been utilized to create a BLAST database to predict epitopes [10].

Machine-learning (ML) approaches have become increasingly important in B-cell epitope prediction [11– 16]. Rather than relying on specific physicochemical characteristics, these methods can learn combinations of sequence and structure derived features from experimentally characterized epitopes and non-epitopes. More recently, artificial intelligence (AI) has revolutionized protein representation [17, 18] and has also influenced B-cell epitope prediction. BepiPred-3 [14] introduced representations generated by protein language models (PLMs) to predict both linear and conformational B-cell epitopes from protein sequence. PLMs [19] learn patterns from very large corpora of protein sequences and can encode information related to structural and functional properties without requiring task-specific training. Such approaches demonstrate the potential of learned protein representations to improve sequence-based B-cell epitope prediction by capturing biologically relevant information beyond conventional sequence-derived features.

Prediction of conformational B-cell epitopes is more challenging because antibody-binding regions are determined by spatial relationships among surface-exposed residues, which cannot be adequately characterized from sequence information alone. The increasing availability of experimentally determined protein structures and high-quality computational structure prediction [20, 21] has expanded the opportunities for incorporating structural information into B-cell epitope analysis. Consequently, rich structure-based representations were also utilized in epitope prediction task to reduce this challenge. For instance, DiscoTope-3 [13] incorporated the inverse folding structure representations [22] to predict the conformational epitopes. Despite these advances, computational prediction of B-cell epitopes remains difficult. Although tremendous advancement has been observed in general protein interaction prediction [20, 23–25], it has made limited progress in B-cell epitope prediction due to lack of experimentally validated data [6]. In addition, antibody recognition is influenced by numerous factors such as specific antibody recognition site and the exact interaction mechanism, much of which is poorly understood [3]. Furthermore, conformational variability of antigenic regions can alter epitope geometry and accessibility, thereby affecting antibody binding [26]. As a result, epitope prediction methods could benefit from incorporating available experimental evidence, including experimentally determined protein structures and known antibody–antigen interactions involving related proteins from the same family or genus.

Here, we present CARTnet, a novel B-cell epitope prediction framework that employs contact-aware attention to integrate sequence, structural, and physicochemical information, including hydrophobicity and electrostatic properties that are often not explicitly considered in recent prediction methods. Furthermore, we introduce two versions of CARTnet: a *de novo* prediction model that operates without prior epitope information, and a template-guided model that can leverage available empirical evidence, such as experimentally characterized antibody-binding regions from structurally related antigens, to guide epitope prediction. Finally, we have also curated and will release a high-quality structural dataset for B-cell epitope prediction and an external dataset for benchmarking.

## 2 Methods

### 2.1 Input Features and Encoding

To capture complementary sequence and structural information, CARTnet integrates multiple residue-level features. We use structural embeddings from ESM3 [27], a large-scale generative protein language model that learns a unified representation across protein sequence, three-dimensional structure, and functional information, rather than modeling these modalities independently. These learned representations are complemented with multimodal features such as binary intra-chain residue contacts [28] to provide topological awareness of the protein structures, eight-state secondary-structure annotations [29], sequence identity, and physico-chemical properties [30], providing both learned and explicit descriptions of the structural and biochemical properties of each antigen residue.

Given an antigen sequence of *L* residues with its coordinates of *C*_*α*_, *N, C* backbone atoms, i.e., (***x*** = ***x***_seq_ ∈ *A*^*L*^, ***x***_coord_ ℝ^*L×*3*×*3^) , CARTnet first extracts a set of static features: 1) ESM3-derived structural embeddings ***F***_ESM_ ∈ ℝ^*L×*1537^ using ***x***_coord_; 2) multimodal residue features ***F***_res_ ∈ ℝ^*L×*32^, e.g., physico-chemical properties and secondary structure, from both sequence and structural information; and 3) intra-chain structural contacts ***C***_intra_ ∈ {0, 1}^*L×L*^. The detailed breakdown of these features is provided in Table S1. The ESM3 embedding ***F***_ESM_ is further projected into a residue-level latent representation, ***H***_ESM_ = MLP_ESM_(***F***_ESM_) through a learnable MLP block. Note that the alphabet *A* includes 20 canonical amino acids and 1 unknown symbol.

### 2.2 Epitope Template as Prior Information

Besides leveraging the above-mentioned multimodal features for epitope site prediction, CARTnet innovates a more flexible architecture that integrates them through a series of contact-aware attention blocks (*N* =10) [31] to update an initial prediction-logit vector ***z*** ∈ ℝ^*L*^:

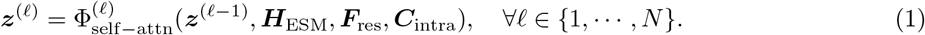

This design choice further enables us to incorporate prior epitope information during prediction when such a template is available. Specifically, this optional template ***m***_temp_ ∈ {0, 1}^*L*^, where 1 indicates a known epitope site, initializes the prediction logit vector ***z***^(0)^. This is updated later to ***z***^(*N*)^ after *N* self-attention blocks with contact map-derived bias, which we discuss next. Note, ***z***^(0)^ = **0** when no template information is available.

### 2.3 Contact-aware Attention ResNet Architecture

Each *ℓ*^th^ contact-aware attention block 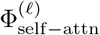 uses the same set of features: (***H***_ESM_, ***F***_res_, ***C***_intra_), to progressively update the prediction logit vector ***z***^(*ℓ*−1)^ to ***z***^(*ℓ*)^. Specifically, it first transforms the current estimate of prediction-logit ***z***^(*ℓ*−1)^ to a *hint* signal 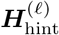 (eq. (2)), which later updates ***H***_ESM_ to ***H***^(*ℓ*)^ (eq. (3)). Regardless of initialization of ***z***^(0)^, the sequential update of ***z*** impacts the successive attention blocks only through this 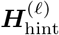:

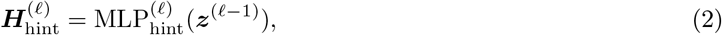

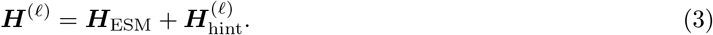

For self-attention of ***H***^(*ℓ*)^, the query and key matrices are obtained from rotary positional embeddings (RoPE) [32] of ***H***^(*ℓ*)^, i.e., ***Q***=***K***=RoPE(***H***^(*ℓ*)^), to incorporate relative sequence positional information. To encourage the model to prioritize structurally interacting residues, we introduce an explicit contact-based bias into the attention mechanism, which we term “contact-aware attention”:

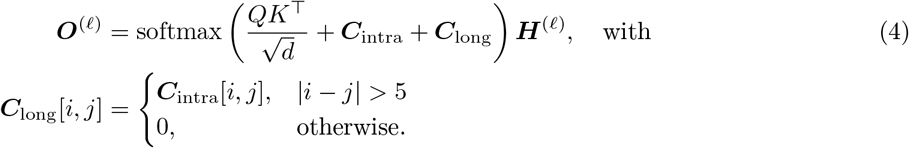

Specifically, the binary intra-chain contact matrix ***C***_intra_ and ***C***_long_ are incorporated into the attention scores where the latter term emphasizes contacts between residues that are distant in the primary sequence. Collectively, this mechanism integrates relative sequence positional information with explicit structural contact information, enabling the attention mechanism to account for both sequential and spatial relationships among residues.

Finally, the resulting attention output ***O***^(*ℓ*)^ is fused with ***H***^(*ℓ*)^ and ***F***_res_ to estimate the logit-scale correction signal, *Δ****z*** = ***z***^(*ℓ*)^ − ***z***^(*ℓ*−1)^. Particularly, the fused embedding ***F***_fused_ is added to ***H***^(*ℓ*)^ in a residual connection manner [33], followed by layer normalization to produce the updated representation ***H***^*′*^:

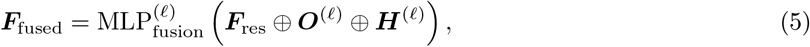

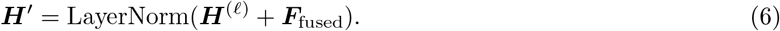

Here, ***H***^*′*^ can be thought of as spatially-aware compressed representation of (***H***_ESM_, ***F***_res_) at each *ℓ*^th^ contactaware self-attention block, conditioned on the estimate of prediction-logit ***z***^(*ℓ*−1)^ from previous block. A prediction MLP then converts it to a residue-level correction *Δ****z***, which is added to the existing signal ***z***^(*ℓ*−1)^:

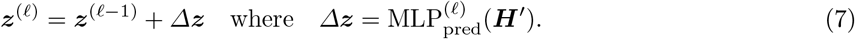

The updated signal ***z***^(*ℓ*)^ is subsequently used to condition the next attention block, enabling information to propagate across successive blocks. After *N* =10 blocks, CARTnet returns the residue-level epitope logits ***z***^(*N*)^ ∈ ℝ^*L*^, which are converted into probabilities using a sigmoid function during evaluation, i.e., ***ŷ*** = *σ*(***z***^(*N*)^). The overall computational procedure is summarized in Algorithm S1 and **Fig. 1** .

**Fig. 1:**
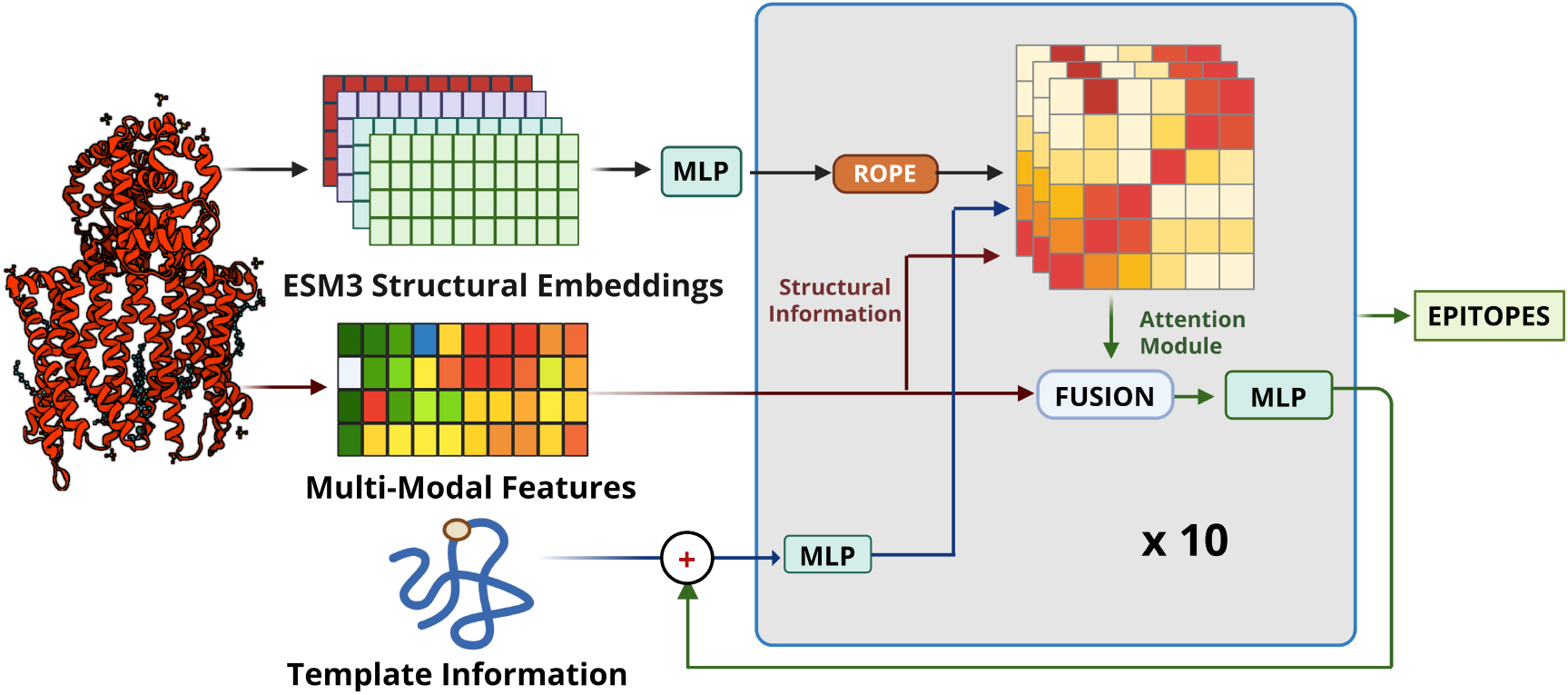
Overview of the CARTnet architecture. ESM3 structural embeddings, multimodal residue features, intra-chain contacts, and optional template information are integrated through 10 contact-aware attention blocks. Structural information biases the attention scores, while the template-guided representation iteratively updates the prediction through residual connections to generate the residue-level epitope predictions.

### 2.4 Training Procedure and Hyperparameters

For an antigen sample ***x***^(*i*)^ with epitope annotations ***y***^(*i*)^ ∈ {0, 1}^*L*^, and the predicted epitope probabilities ***ŷ***^(*i*)^ ∈ [0, 1]^*L*^, the training objective of CARTnet integrates a focal loss [34], a target-wise Dice loss [35], and a template neighborhood loss:

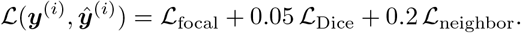

The focal loss *ℒ* _focal_ (see section S1.3 for details of the loss components) is applied with *γ* = 2 and *α* = 0.7 to address the imbalance between epitope and non-epitope residues, while ℒ_Dice_ encourages overlap between predicted and true epitope residues. The template-neighborhood loss applies a masked focal loss on non-template residues that are in direct structural contact (based on ***C***_intra_) with the provided template residues in ***m***_temp_. Since the template residues themselves are excluded from this auxiliary loss, ℒ_neighbor_ encourages the model to selectively identify additional epitope residues in the vicinity of the template, rather than indiscriminately assigning high epitope scores to all residues near the templates. When no template is used or selected, ℒ_neighbor_ = 0.

The network is optimized using AdamW [36, 37] with a learning rate of 5 *×* 10^−5^, weight decay of 1 *×* 10^−5^, and a batch size of 16. Model performance is monitored on the validation set during training, and the model achieving the highest validation area under the precision–recall curve (AUPRC) is selected for final evaluation. During training of CARTnet with template information, a small number of true labels are randomly selected as templates [38], as detailed in Section S1.4 of the supplementary document.

### 2.5 Datasets and Evaluation Metrics

The dataset has been curated from the Antigen-Antibody Complex Database (AACDB) [39], which contains experimentally characterized antigen and antibody complexes released till November 2023 in the Protein Data Bank (PDB) [40]. Altogether, the database comprises 7,498 high-quality, manually curated antigen–antibody complexes. The data from AACDB have been filtered to remove any sequence less than or equal to 15 residue length or greater than 1400. In addition, the structures have also been cleaned to remove various inconsistencies. This includes removing duplicate *C*_*α*_ atom records, resolving alternate atom locations, removing nonstandard residues, and standardizing residue numbering.

CD-HIT [41] has been used to reduce sequence redundancy during dataset construction. A 50% sequence identity threshold has been first applied to the complete dataset, and cluster representatives have been selected as targets for the training set, following a strategy similar to previous related works [13, 14]. After the training set is separated, a more stringent 40% sequence identity threshold is applied between the training set and the validation and internal test sets to further reduce sequence similarity across the data splits to avoid potential data leakage as much as possible. Detailed statistics for these datasets are provided in Supplementary Table S2.

In our experiments, we have defined epitope as a residue on an antigen recognized and bound by an antibody. For structure-based annotation, different distance thresholds have been used to identify epitope residues ranging from 4 Å to 6 Å [13, 14, 39, 42–44]. To maintain consistency with the AACDB dataset used in this study, we define an antigen residue as an epitope residue if it contains at least one heavy atom within 6 Å of an antibody residue [39, 44]. Furthermore, epitope collapsing has been performed at 95% sequence identity [14], whereby epitope annotations from highly similar antigen sequences are transferred to the corresponding positions of the cluster representative, thereby increasing the availability of positive epitope labels.

In addition to this curated internal test set, an external benchmark test set has been constructed to evaluate generalization to substantially different antigens. The external test set includes antigen–antibody complexes consisting of structures from the Structural Antibody Database (SAbDab) [45] as well as bench-mark antigens used in previous B-cell epitope prediction studies [13, 14]. We have collected complexes from SAbDab with release dates after January 1, 2025, retained only entries classified as Fv, and applied the same filtering criteria used for the previously described dataset. Targets from the external sequences are further screened against the training data and themselves, to remove any sequences exceeding 20% sequence identity. We have also removed complexes that have binding partners other than antibody. This stringent similarity filtering is used to provide a more challenging assessment of model performance on antigens with limited sequence similarity to those encountered during training. It is noteworthy that no epitope collapsing is implemented for this dataset.

Model performance has been evaluated using both target-wise and global metrics. For each antigen target, precision, recall, F1-score, accuracy, and Matthews correlation coefficient (MCC) are calculated individually and then averaged across all targets for their top-20% prediction. We also calculate the precision at Top-10 using the top 10 predictions. In addition, predictions from all residues across the dataset are pooled to calculate global area under the receiver operating characteristic curve (AUROC) and area under the precision–recall curve (AUPRC). All metrics, except Precision at Top-10, are calculated using scikit-learn [46]. We use the top-*k*% ranked residue predictions to calculate the evaluation metrics, thereby avoiding inconsistencies arising from the different epitope-classification thresholds used by individual prediction tools.

## 3 Results

### 3.1 Performance on the Internal Test Set

The CARTnet models (with and without template) were evaluated on the internal test set comprising 61 targets and compared with BepiPred-2 [16], BepiPred-3 (BP3) [14] and DiscoTope-3 (DT3) [13] (**Table 1** and **Fig. S1**). Both CARTnet variants were evaluated without supplied template residues in these benchmark comparisons; CARTnet_W_temp denotes the model trained to accept optional template information. To replicate non-ideal scenarios where no experimentally validated structure would be present, we evaluated with ColabFold [47] predicted structures also for both DT3 and CARTnet. Thus, for structure-based configurations, *true* denotes inference using experimentally determined structures, whereas *af2* denotes inference using predicted ColabFold structures [47].

**Table 1:** Performance comparison of different models and configurations on the internal test set consisting of 61 targets. Best-performing values are highlighted in **bold**.

| **Model** | **Accuracy** | **Precision** | **Recall** | **F1** | **AUC** | **AUPRC** | **Prec@Top-10** | **MCC** |
|---|---|---|---|---|---|---|---|---|
| CARTnet_W/O_temp_true | **0.754** | **0.301** | **0.411** | **0.315** | **0.700** | **0.243** | 0.344 | **0.196** |
| CARTnet_W/O_temp_af2 | 0.733 | 0.249 | 0.339 | 0.260 | 0.674 | 0.219 | 0.266 | 0.124 |
| CARTnet_W_temp_true | 0.749 | 0.289 | 0.396 | 0.303 | 0.677 | 0.225 | **0.352** | 0.181 |
| CARTnet_W_temp_af2 | 0.731 | 0.244 | 0.330 | 0.254 | 0.654 | 0.200 | 0.270 | 0.116 |
| DiscoTope-3_true | 0.740 | 0.266 | 0.346 | 0.272 | 0.675 | 0.220 | 0.318 | 0.142 |
| DiscoTope-3_af2 | 0.735 | 0.255 | 0.320 | 0.260 | 0.651 | 0.195 | 0.274 | 0.123 |
| BepiPred-3 | 0.722 | 0.224 | 0.294 | 0.235 | 0.651 | 0.198 | 0.220 | 0.086 |
| BepiPred-2 | 0.714 | 0.202 | 0.239 | 0.200 | 0.531 | 0.140 | 0.205 | 0.045 |

Among all evaluated methods, CARTnet_W/O_temp_true achieved the best overall performance. Compared with DiscoTope-3_true, the strongest baseline across most metrics, CARTnet improved F1 from 0.272 to 0.315, AUC from 0.675 to 0.700, and AUPRC from 0.220 to 0.243. Notably, the improvement in F1 resulted from simultaneous increases in both precision (0.301 vs. 0.266) and recall (0.411 vs. 0.346), indicating an improved balance between sensitivity and precision.

The proposed models also performed favorably in identifying highly ranked epitope residues. CART-net_W_temp_true achieved the highest Top-10 precision (0.352). Overall, configurations using experimentally determined structures consistently outperformed their predicted-structure counterparts, although the latter remained competitive with the baseline methods.

Additionally, using predicted structure as input (CARTnet_W/O_temp_af2) performed better than DiscoTope-3_af2 in the exact setup by most metrics except for precision-based metrics. For instance, CART-net_W/O_temp_af2 achieved an AUPRC of 0.219 in comparison to 0.195 of DiscoTope-3_af2.

### 3.2 Performance on the External Test Set

Generalization was further evaluated on an independent external test set comprising 52 targets (**Table 2**). CARTnet_W/O_temp_true achieved the strongest overall performance, with an accuracy of 0.767, precision of 0.275, recall of 0.420, F1 score of 0.298, AUC of 0.743, AUPRC of 0.267, and MCC of 0.196.

**Table 2:**
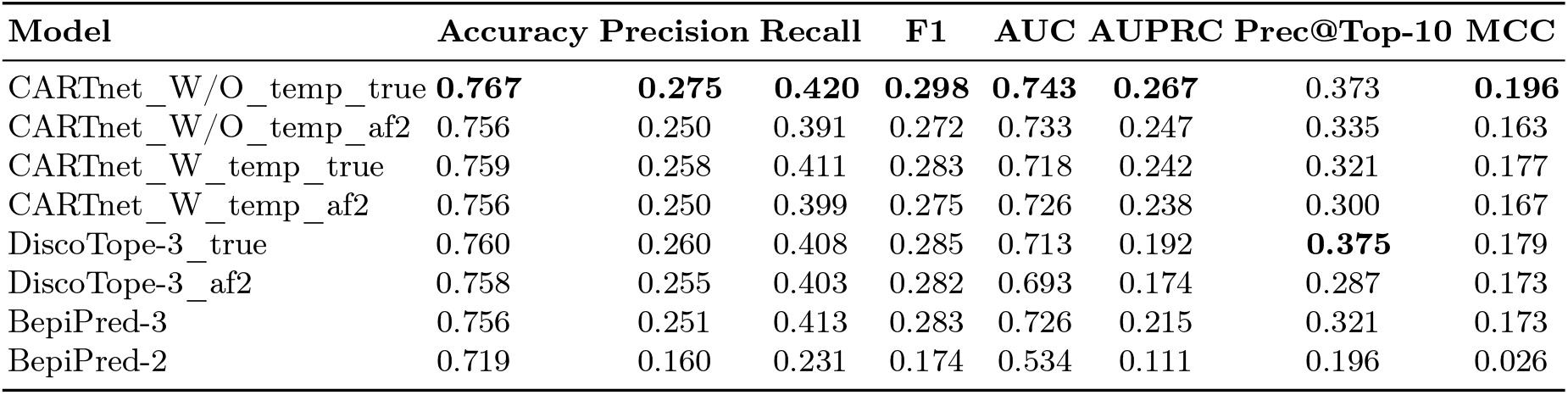
Performance comparison of different models and configurations on the external test set of 52 targets. Best-performing values are highlighted in **bold**.

The greatest improvement over the established predictors was observed in AUPRC. For example, CART-net_W/O_temp_true achieved an AUPRC of 0.267, compared with 0.215 for BepiPred-3 and 0.192 for DiscoTope-3_true, corresponding to absolute improvements of 0.052 and 0.075, respectively. These represent relative improvements of 24.2% over BepiPred-3 and 39.1% over DiscoTope-3_true. The improvement in AUPRC was accompanied by higher precision and recall: CARTnet_W/O_temp_true achieved a precision of 0.275 and recall of 0.420, compared with 0.251 and 0.413 for BepiPred-3 and 0.260 and 0.408 for DiscoTope-3_true, respectively. Consequently, the proposed model also achieved the highest F1 score and MCC on the external benchmark.

For the top-ranked predictions, CARTnet_W/O_temp_true achieved a Top-10 precision of 0.373, compared with 0.321 for BepiPred-3 and 0.375 for DiscoTope-3_true. Although DiscoTope-3_true marginally outperformed the proposed model in Top-10 precision, the proposed model demonstrated stronger performance across a broader range of classification and ranking metrics.

It is noteworthy that the performance of the variants of CARTnet was generally higher when experimentally determined structures were used instead of predicted structures. Nevertheless, the predicted-structure configurations retained competitive performance, demonstrating the applicability of the proposed framework when experimentally determined structures are unavailable.

### 3.3 Impact and Robustness of Template-guided Prediction

To evaluate whether prior knowledge of an epitope can improve the identification of additional epitope residues on the same antigen, we performed a controlled template-guided prediction experiment on the internal test set. For each target antigen, a fixed number of experimentally annotated epitope residues were randomly selected and provided to CARTnet_W_temp as template information during inference. The remaining epitope annotations were retained for evaluating its predictions, i.e., provided epitope residues were removed from the evaluation. The number of randomly selected templates was varied from 0 to 9. The performance evaluation results are presented in **Table S6** and **Fig. 2** left panel. The trend clearly shows that increasing the number of templates increased the performance of the CARTnet. The AUPRC of the CARTnet_W_temp was 0.225 and after including only 1 template epitope it rose to 0.403. However, it is also clear that the rate of increase in performance diminished when increasing the number of template epitopes and eventually plateaued. This saturation may be attributed to the finite number of epitope residues available for identification, which limits further performance gains.

**Fig. 2:**
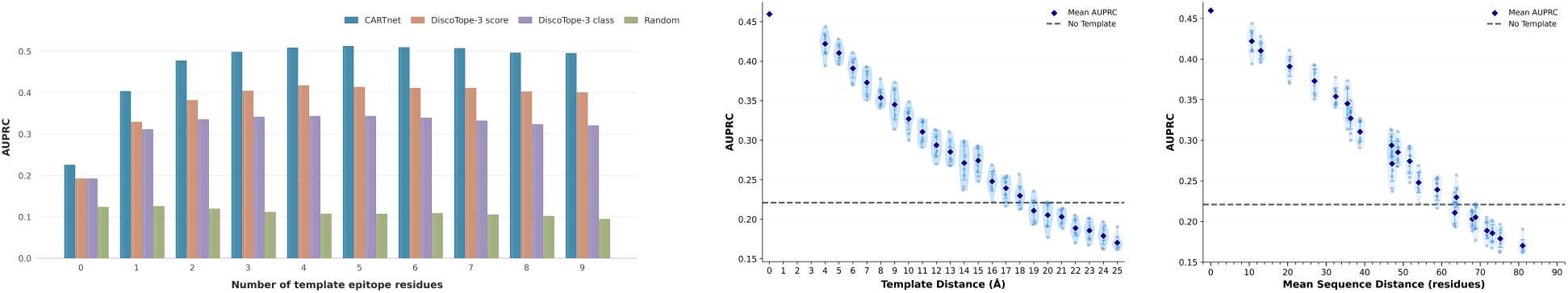
Template-guided prediction analysis. The left panel displays the impact of increasing the number of template epitope residues on the AUPRC. The middle and right panels display the effect of perturbation on the AUPRC in terms of spatial distance and sequence distance, respectively.

**Fig. 3:**
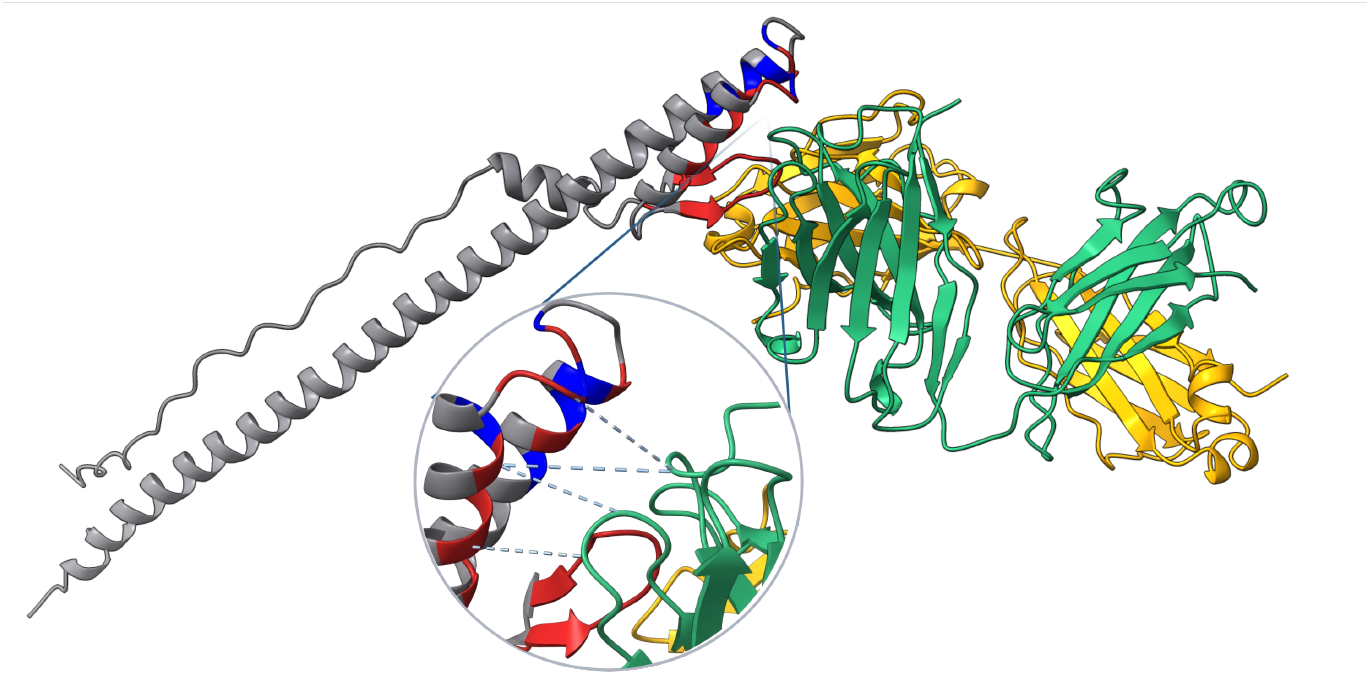
Illustration of the S1V2-72 Fab–EHA2 complex from influenza B/Malaysia/2506/2004. The antibody chains are shown in green and yellow, and the antigen chain is shown in gray. Experimentally annotated epitope residues are shown in blue, and those correctly predicted by CARTnet are shown in red.

In addition, we evaluated two simple template-guided baselines using DiscoTope-3. DiscoTope-3 residue scores were first min–max normalized to the range [0,1]. For DiscoTope-3 score, template information was incorporated by adding 0.5 to residues in contact with a template epitope residue, an additional 0.5 if the residue itself corresponded to the template epitope, and 0.25 *×* RSA, where RSA denotes relative solvent accessibility. This scoring scheme therefore strongly prioritizes template residues and their spatially neigh-boring surface-exposed residues while retaining the original DiscoTope-3 score. For the DiscoTope-3 class, the normalized DiscoTope-3 scores were retained for all residues, except that residues in contact with a template epitope residue and having RSA *>* 0.2 were assigned the maximum normalized score of 1.0. All remaining residue scores were left unchanged. The resulting residue-level scores were then used directly to calculate the global AUPRC. Both approaches performed substantially worse than CARTnet, suggesting that simple rule-based incorporation of template information is less effective than using a model specifically trained to integrate template-derived information.

To determine how sensitive CARTnet is to inaccuracies in the supplied template, we performed a controlled template-perturbation experiment. For each target in the internal test set, one experimentally annotated epitope residue was selected as the true template at random. We then introduced spatial errors into the template by replacing the true template residue with another residue located at a predefined distance from it in the three-dimensional structure. For a target perturbation distance *d*, candidate residues were selected within the interval *d* − 4 to *d* Å from the true template residue. The perturbation distance was progressively increased from 4 Å to 25 Å, allowing us to evaluate how prediction performance changes as the supplied template becomes increasingly displaced from the experimentally identified epitope. Since multiple residues can satisfy a given distance criterion, the perturbation procedure was repeated 10 times at each distance, with candidate erroneous template residues sampled independently for each replicate. For each run, AUPRC was calculated to quantify the effect of increasing template displacement on prediction performance. Since only one epitope residue was used, it was not removed during evaluation. One target (PDB ID: 4ZFO) was excluded from this analysis because its short sequence did not contain sufficient candidate residues across the required perturbation-distance ranges.

The results demonstrate that CARTnet is sensitive to the spatial accuracy of the supplied template, with AUPRC progressively decreasing as the template was displaced from the true epitope residue (**Fig. 2**-middle panel). However, the decline in performance was gradual rather than abrupt. AUPRC decreased from approximately 0.46 with the correct template to 0.35 at an 8 Å perturbation and remained above the no-template baseline (AUPRC *∼* 0.22) for perturbations extending to approximately 18 Å. At larger perturbation distances, the benefit of the template was lost, with performance eventually falling below the no-template baseline. These results indicate that although CARTnet benefits from spatially accurate template information, it can tolerate substantial spatial deviations before the template ceases to provide useful information.

We further examined the sequence separation between the true and perturbed template residues (**Fig. 2**-right panel). AUPRC generally decreased as the sequence distance increased, although the relationship was less consistent than for spatial distance. Nevertheless, CARTnet could tolerate a sequence separation of 60 residues. However, a large sequence separation does not necessarily imply a large spatial separation in the folded protein. These results strongly indicated that CARTnet tolerates moderate template errors, whereas increasingly inaccurate templates gradually reduce their benefit and can eventually perform worse than using no template.

### 3.4 Case Study: Improved B-Cell Epitope Prediction in Influenza Virus

Influenza viruses cause recurrent seasonal epidemics worldwide and remain important targets for vaccine and therapeutic antibody development. The continuous antigenic evolution of influenza viruses makes the identification and characterization of antibody-binding epitopes particularly important. To investigate the performance of CARTnet_W/O_temp on a biologically relevant viral antigen, we selected the S1V2-72 Fab–EHA2 influenza complex (PDB ID: 8UDG) from our internal test set as a case study. The experimentally resolved antibody–antigen complex provides well-defined epitope residues, allowing residue-level comparison between the experimentally observed epitope and the predictions generated by CARTnet.

CARTnet showed strong performance on this target, successfully identifying a substantial proportion of the experimentally determined epitope residues, as shown in **Table S3**. Importantly, the predicted residues were predominantly localized around the experimentally observed antibody-binding region. CART-net achieved a recall of 0.76, substantially higher than those of DiscoTope-3 (0.440), BepiPred-3 (0.200), and BepiPred-2 (0.000), demonstrating its ability to identify the spatial pattern of the experimentally characterized epitope on the antigen surface.

### 3.5 Case Study 2: Generalizing Epitope Information Within the Genus Alphavirus

Alphaviruses are mosquito-borne viruses that include several important human pathogens, including Chikungunya virus (CHIKV), Mayaro virus (MAYV), and Ross River virus (RRV). These related viruses share similarities in their structural proteins while causing distinct outbreaks and disease burdens. When experimentally characterized epitope information is available for one member of such a related viral group, this information may provide a useful starting point for investigating antibody-binding regions in other members for which experimental information is limited. To investigate this application of CARTnet, we used experimentally characterized epitope information from Ross River virus (PDB ID: 6VYV) and extracted its epitope as a template to guide epitope prediction for Chikungunya and Mayaro viruses, illustrated in **Fig. 4. Table S4** summarizes the quality of the transferred epitope templates. The target antigens share 56.6–57.0% sequence identity with the RRV sequence by BLAST [48], while their structures show high similarity to the RRV structure, with TM-scores ranging from 0.907 to 0.917 measured using MM-align [49]. The experimentally characterized RRV epitope residues were first mapped onto the RRV sequence and subsequently transferred to the corresponding target residues through sequence alignment to generate the template information provided to CARTnet. We further evaluated the quality of the transferred templates by comparing the mapped epitope residues with the experimentally determined epitope labels of MAYV (PDB ID: 6W1C) and CHIKV (PDB ID: 6W09). The transferred templates recovered 83.3% and 61.1% of the experimentally characterized epitope residues in MAYV and CHIKV, respectively, indicating that the template information was informative but incomplete.

**Fig. 4:**
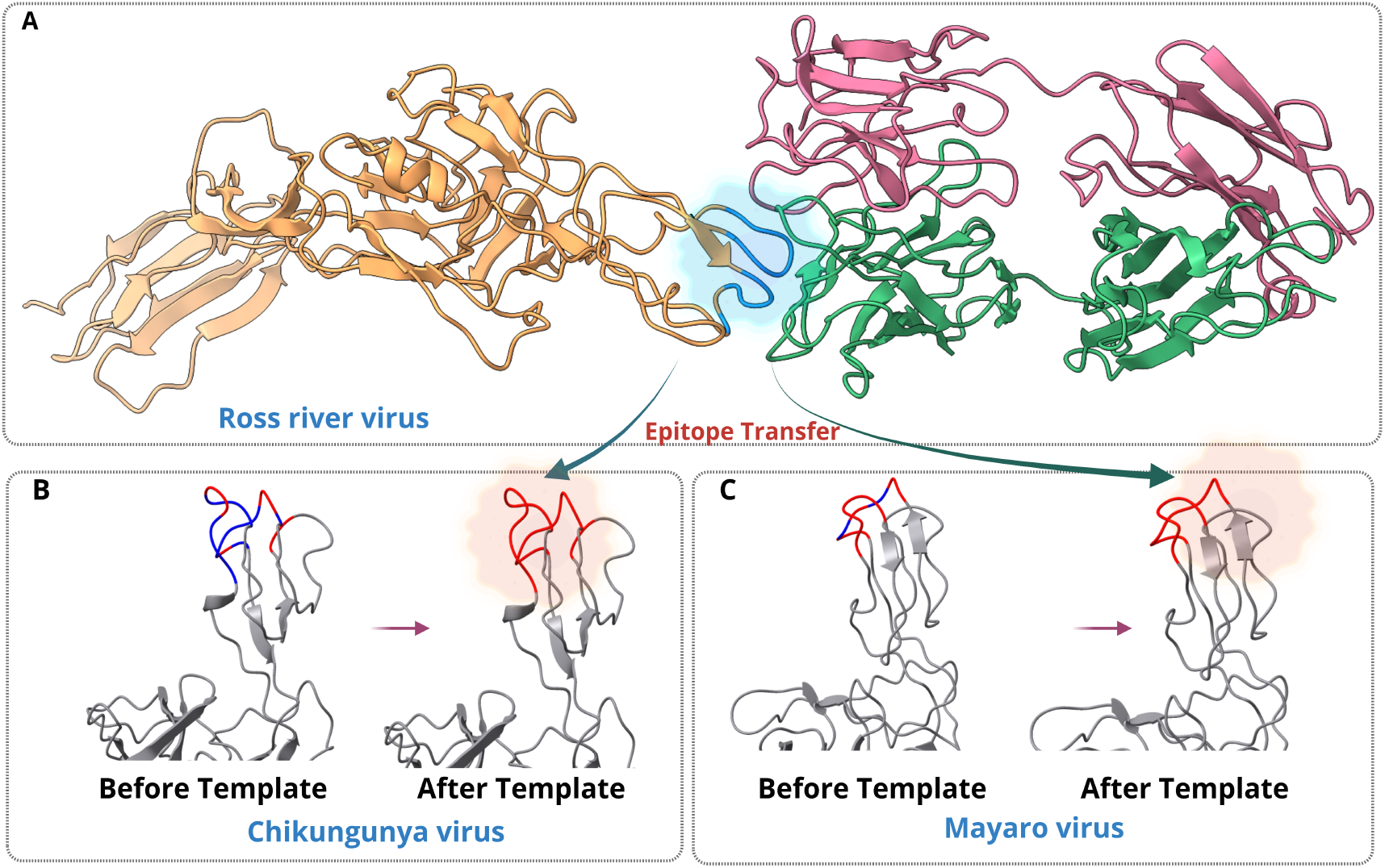
Illustration of the epitope transfer mechanism of CARTnet. (A) The experimentally characterized epitope is extracted from RRV and mapped to the corresponding regions of CHIKV and MAYV to provide template information. (B, C) Epitope predictions for CHIKV and MAYV, respectively, before and after incorporating the transferred template information. Blue indicates experimentally characterized ground-truth epitope residues, while red indicates correctly predicted epitope residues. After incorporating the transferred epitope information, CARTnet successfully identified all ground-truth epitope residues in both targets.

Incorporating RRV epitope information as a template substantially improved predictions for both related viruses (**Table 3**). For this case study, the transferred template residues were retained during evaluation. For CHIKV (6W09), recall increased from 0.389 to 1.000, while for MAYV (6W1C), recall increased from 0.833 to 1.000. Furthermore, Prec@Top-10 increased from 0.200 to 1.000 for CHIKV and from 0.200 to 0.900 for MAYV, demonstrating that template information substantially improved the ranking of true epitope residues among the highest-confidence predictions. Precision and F1-score also improved for both targets. Notably, although the transferred templates themselves recovered only 61.1% and 83.3% of the experimentally characterized CHIKV and MAYV epitopes, respectively, CARTnet achieved 100% epitope recall after incorporating this partial information. This demonstrates the ability of CARTnet to use incomplete epitope information from a related viral antigen to identify additional epitope residues in the target antigen.

**Table 3:**
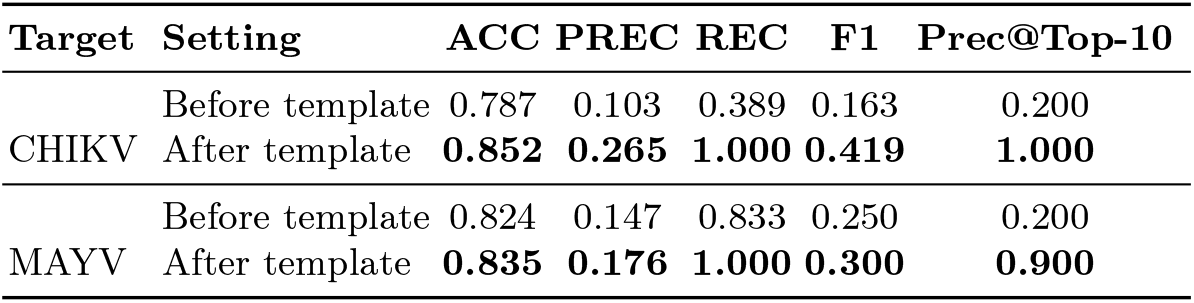
Effect of template information on the performance in the Alphavirus case-study. Performance before and after incorporating template information for CHIKV (6W09) and MAYV (6W1C).

| **Target** | **Setting** | **ACC** | **PREC** | **REC** | **F1** | **Prec@Top-10** |
|---|---|---|---|---|---|---|
| CHIKV | Before template | 0.787 | 0.103 | 0.389 | 0.163 | 0.200 |
|  | After template | **0.852** | **0.265** | **1.000** | **0.419** | **1.000** |
| MAYV | Before template | 0.824 | 0.147 | 0.833 | 0.250 | 0.200 |
|  | After template | **0.835** | **0.176** | **1.000** | **0.300** | **0.900** |

## 4 Discussion and Conclusion

In this study, we have developed CARTnet, a novel contact-aware deep learning framework for residue-level B-cell epitope prediction that integrates multimodal features and optional template-derived epitope information. CARTnet consistently outperforms the evaluated sequence and structure-based baselines on both the curated internal test set and the stringent external test set. The preservation of this performance advantage on the external dataset highlights CARTnet’s ability to generalize to antigens substantially different from those encountered during training. In addition to that, it is also observed that CARTnet also performs better when an experimental structure is available, though performance with predicted structures remains comparable.

Learned structural embeddings capture complex residue-level patterns, while explicit physicochemical features provide complementary biochemical information that may not be explicitly represented in these embeddings. Intra-chain contacts further capture spatial relationships between residues that are distant in sequence but proximal in three-dimensional space, a key characteristic of conformational B-cell epitopes. By integrating these complementary features through contact-aware attention, CARTnet can achieve improved epitope prediction performance, as supported by the benchmark and ablation results.

The ablation experiments presented in Supplementary Information S3.2 indicate that different information sources incorporated into CARTnet provide complementary contributions. The largest reduction in performance occurred after removing the ESM3 structural embeddings, resulting in a 31.11% decrease in F1-score, whereas removal of intra-chain contacts or replacement of the contact-aware attention module produced smaller but measurable reductions. These observations suggest that learned structural representations provide the strongest individual contribution, while explicit structural contacts and the specialized attention mechanism further enhance predictive performance. In particular, incorporating intra-chain contacts directly into the attention calculation enables the model to explicitly capture the topological relationships between residues.

Beyond conventional *de novo* epitope prediction, a novel feature of CARTnet is its ability to incorporate previously characterized epitope information to guide the identification of additional epitope residues. This enables the framework to leverage existing experimental knowledge rather than relying exclusively on sequence and structural features, potentially improving prediction reliability for challenging antigen targets. Providing increasing amounts of experimentally annotated template information substantially improves prediction performance, with AUPRC increasing from 0.225 without template information to 0.403 after providing only a single template residue and reaching approximately 0.50 with several template residues. The template-perturbation analysis further demonstrates that this benefit is not restricted to perfectly accurate template information. Thus, CARTnet appears capable of tolerating moderate noise in transferred epitope information, although sufficiently inaccurate templates can eventually become detrimental. This suggests that even limited prior knowledge of an epitope region can guide the model toward additional epitope residues rather than simply reproducing the supplied template.

Furthermore, the genus Alphavirus case study illustrates a potential practical application of the templateguided formulation, demonstrating CARTnet’s ability to transfer partial epitope information from related antigens to identify the complete epitope region, achieving a recall of 1.0. Similarly, the influenza case study demonstrates CARTnet’s effectiveness in a prediction setting without prior epitope information, achieving a recall of 0.76 and a precision of 0.594. Together, these case studies highlight the potential applicability of CARTnet to both template-guided epitope identification and epitope prediction without prior epitope information.

Several limitations should nevertheless be considered. First, CARTnet remains dependent on the quality of the available structural representation. Across both evaluation datasets, configurations using experimentally determined structures generally performed better than those using predicted structures, though the predicted-structure variants remained competitive with existing approaches. Second, B-cell epitope annotation is inherently incomplete because experimentally characterized antibody-antigen complexes represent only a subset of possible antibodies, antigen conformations, and accessible binding sites. Finally, although template-guided prediction showed strong improvements in controlled experiments and in the Alphavirus case study, broader evaluation across additional antigen families will be required to determine how template usefulness depends on sequence similarity, structural similarity, and epitope conservation.

Overall, CARTnet demonstrates the effectiveness of integrating learned structural representations, explicit residue-level features, and contact-aware attention for B-cell epitope prediction. Beyond this, CARTnet extends conventional *de novo* prediction by allowing experimental knowledge from related antigens to be incorporated directly into the prediction process. This complementary approach reuses previously characterized epitope information to refine predictions and make better use of existing experimental data.

## Acknowledgements

This work was supported by the Advanced Research Projects Agency for Health (ARPA-H) Award 1AY1AX000053-01.

## Supplementary Information

## S1 Method and Materials Details

### S1.1 Detailed Feature Description

**Table S1:** Composition and dimensionality of CARTnet input features.

| Feature Block | Component | Dimension |
|---|---|---|
| *F*<sub>ESM</sub> | ESM3 structural embedding | *L* × 1536 |
|  | Sequence-length indicator (binary) | *L* × 1 |
|  | **Final concatenated features** | **L × 1537** |
| *F*<sub>res</sub> | Sequence identity (one-hot) | *L* × 21 |
|  | Secondary structure (DSSP) | *L* × 8 |
|  | Charge | *L* × 1 |
|  | Hydrophobicity | *L* × 1 |
|  | Sequence-length indicator (binary) | *L* × 1 |
|  | **Final concatenated residue features** | **L × 32** |
| *C*<sub>intra</sub> | Binary intra-chain contact map | *L* × *L* |

### S1.2 CARTnet Algorithm

#### Algorithm S1

CARTnet: Contact-Aware Residual Attention for B-cell Epitope Prediction

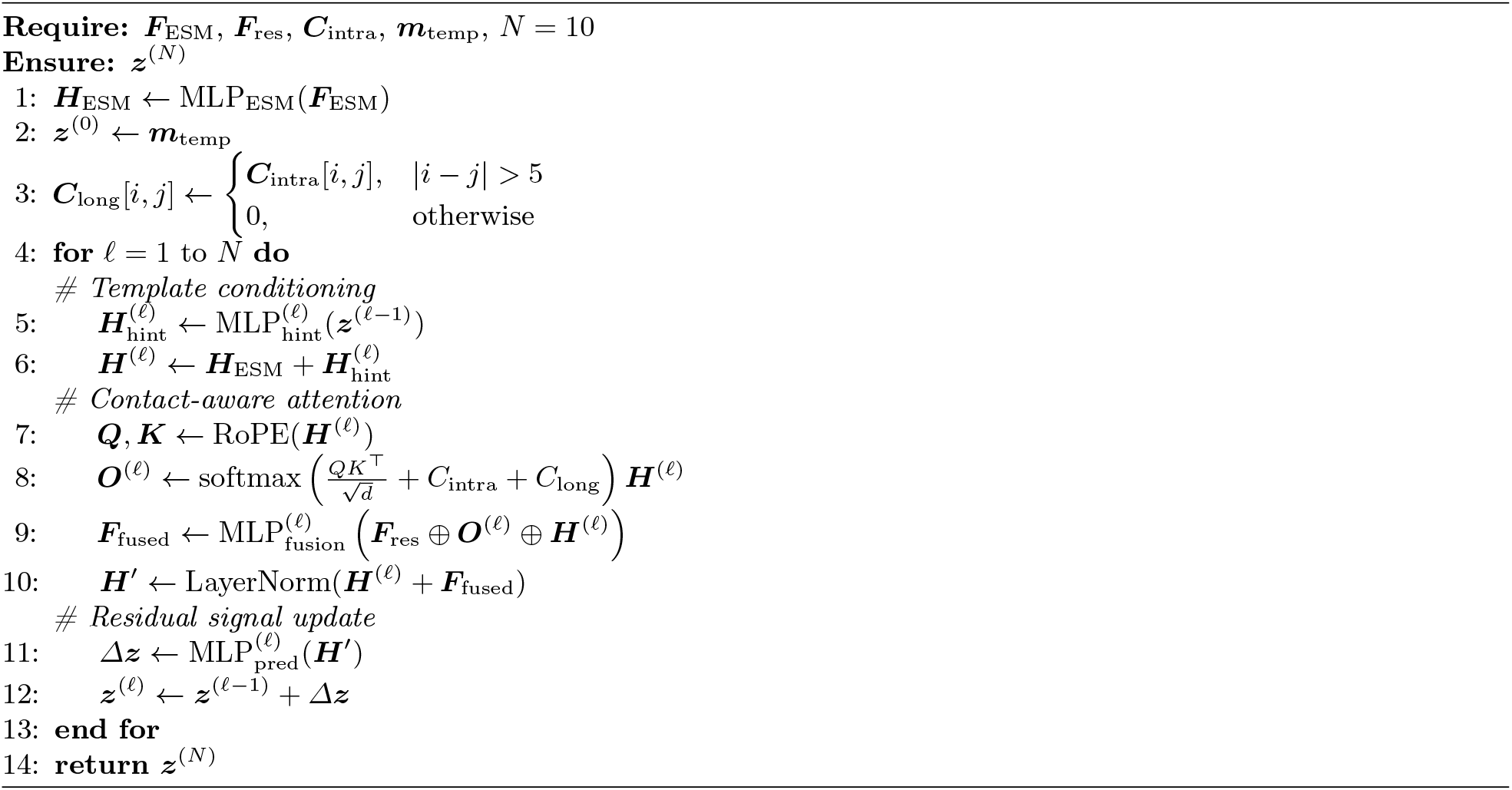

### S1.3 Details of Loss Components

Given the pair of epitope annotations ***y*** ∈ {0, 1}^*L*^, and the prediction of epitope probability ***ŷ*** ∈ [0, 1]^*L*^ for each sample, the three losses are computed as follows:

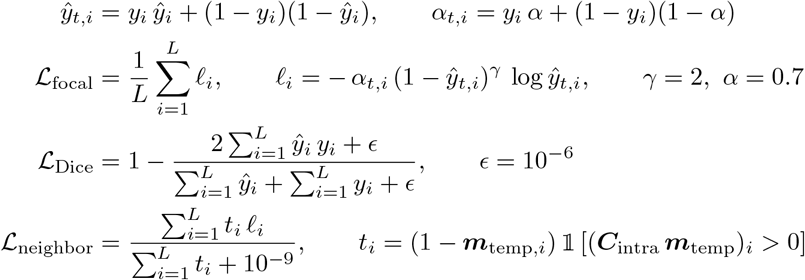

### S1.4 Randomized Template Residue Sampling

During training, template residues were randomly selected from the ground-truth epitope residues. First, the number of template residues (k) was determined through weighted sampling from a predefined set of candidate counts: 0, 1, 2, 3, 4, 10%, or 20% of the total positive residues. Each candidate was assigned a corresponding sampling weight of 0.1, 0.1, 0.1, 0.05, 0.05, 0.1, and 0.1, respectively. Based on the sampled count (k), the corresponding number of positive residue indices was randomly selected without replacement to form the template set (S). This strategy allowed the model to be trained with varying amounts of available template information, including cases where no template residues were provided.

#### Algorithm S2

Randomized epitope template sampling. *L* is the antigen sequence length, *P* is the set of positive residue indices, and *K* and *W* are the candidate template counts and corresponding weights. Weighted sampling determines the number of templates *k*, and random sampling selects their indices *S*.

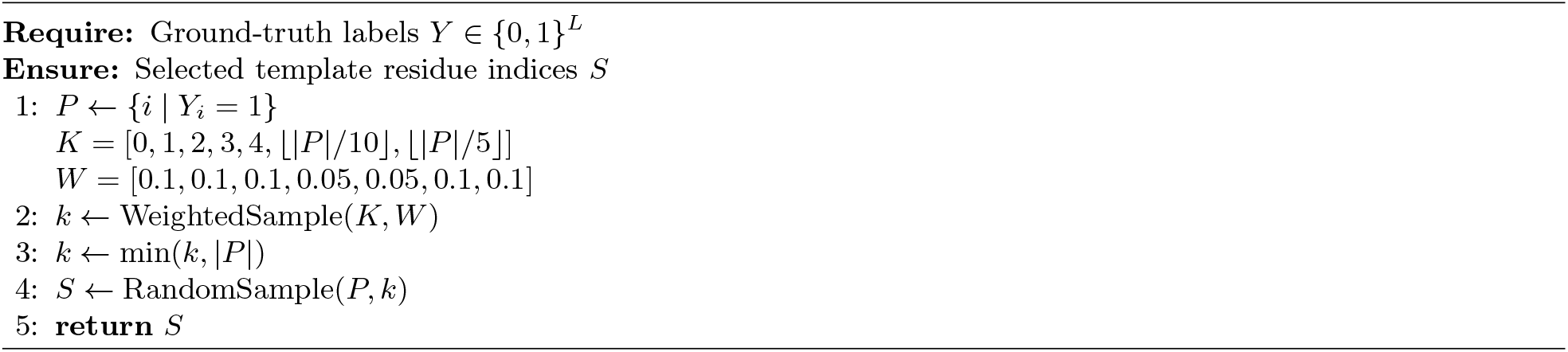

### S1.5 Dataset Statistics

We recorded the pLDDT confidence score reported by ColabFold for each predicted antigen structure in the internal test and external test sets. TM-scores were calculated using MM-align by comparing each predicted antigen structure with its corresponding experimentally determined structure. Both metrics were then averaged across structures separately for each test set. These metrics were not calculated for the training and validation sets because predicted structures were not used for those sets.

**Table S2:** Dataset bstatistics. Summary statistics of dataset size, epitope density, and sequence length across datasets. Mean pLDDT and TM-score values are also reported for predicted structures in the test and external test sets.

|  |  | **Epitope Density** |  |  | **Sequence Length** |  |  | **Predicted Structure Quality** |  |
|---|---|---|---|---|---|---|---|---|---|
| **Dataset** | **Size** | **Min** | **Mean** | **Max** | **Min** | **Mean** | **Max** | **pLDDT** | **TM-score** |
| Training | 542 | 0.017 | 0.189 | 0.889 | 18 | 282.98 | 1265 | - | - |
| Validation | 55 | 0.023 | 0.200 | 0.728 | 43 | 309.91 | 1390 | - | - |
| Internal test set | 61 | 0.029 | 0.166 | 0.559 | 36 | 304.38 | 1267 | 87.442 | 0.831 |
| External test set | 52 | 0.023 | 0.143 | 0.511 | 47 | 287.83 | 1277 | 89.966 | 0.865 |

## S2 Supplementary Performance Analysis

### S2.1 Precision–Recall Curve Analysis

**Fig. S1:**
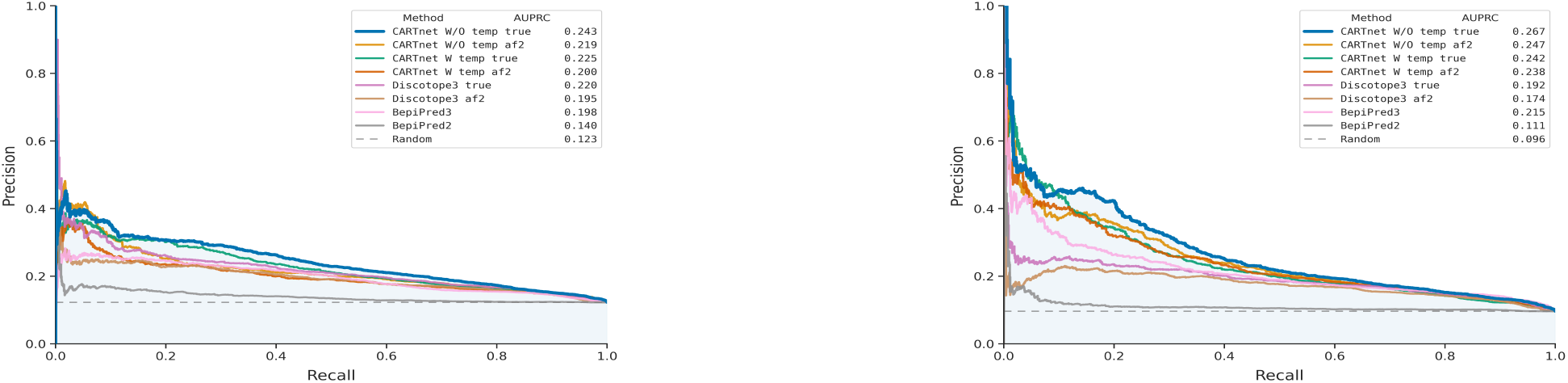
Precision–recall curves comparing CARTnet and baseline methods on the internal (left) and external (right) test sets. The curves illustrate the precision–recall trade-offs across classification thresholds.

### S2.2 Case Study : Improved B-Cell Epitope Prediction in Influenza Virus

**Table S3:** Performance comparison of different models on S1V2-72 Fab bound to EHA2 from influenza B/Malaysia/2506/2004. Best-performing values are highlighted in bold.

| **Model** | **Accuracy** | **Precision** | **Recall** | **F1** | **Prec@Top-10** | **MCC** |
|---|---|---|---|---|---|---|
| CARTnet_W/O_temp | **0.881** | **0.594** | **0.760** | **0.667** | **0.800** | **0.602** |
| DiscoTope-3 | 0.780 | 0.344 | 0.440 | 0.386 | 0.300 | 0.257 |
| BepiPred-3 | 0.704 | 0.156 | 0.200 | 0.175 | 0.200 | -0.001 |
| BepiPred-2 | 0.642 | 0.000 | 0.000 | 0.000 | 0.000 | -0.217 |

### S2.3 Case Study 2 :Generalizing Epitope Information Within the Genus Alphavirus

To assess the quality of the aligned epitope information used as a template, we compared the binary template labels directly with the experimentally determined epitope labels of MAYV and CHIKV (**Table S4**). Since both sets of labels were binary, accuracy, precision, recall, and F1-score were calculated directly.

**Table S4:** Quality of epitope templates used in the case studies of MAYV (6W1C) and CHIKV (6W09). Template–target similarity was assessed using sequence identity and TM-score. The quality of the transferred epitope information was evaluated by directly comparing the binary template labels with the experimentally defined epitope labels of the target antigen.

| **Target** | **Seq. Identity (%)** | **TM-score** | **Accuracy** | **Precision** | **Recall** | **F1** |
|---|---|---|---|---|---|---|
| MAYV | 56.60 | 0.917 | 0.991 | 0.909 | 0.833 | 0.870 |
| CHIKV | 57.00 | 0.907 | 0.979 | 1.000 | 0.611 | 0.759 |

## S3 Additional Analysis of Factors Affecting Model Performance

### S3.1 Impact of Epitope Density on Performance

**Fig. S2:**
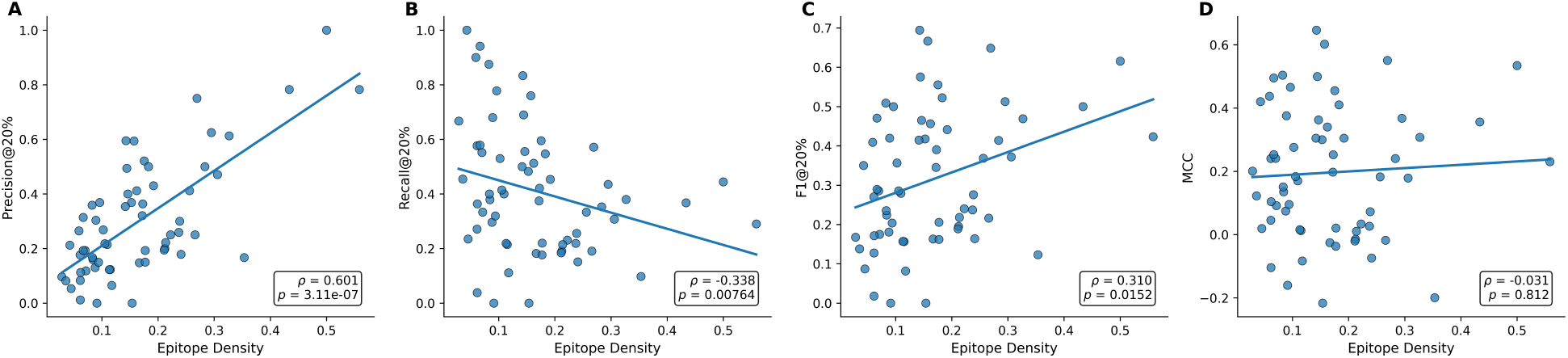
Showing the impact of epitope density on precision, recall, F1-score, and MCC on the test set

An interesting observation in several protein-prediction tasks, such as inter-chain contact prediction, is that predictive performance can depend on the density of positive labels. We therefore investigated whether a similar relationship exists for CARTnet_W/O_temp_true. Epitope density was defined as the fraction of residues annotated as epitopes in each antigen, and its association with per-target precision, recall, F1-score, and MCC was assessed using Spearman’s rank correlation on the top-20% predictions.

As shown in **Figure S2**, epitope density exhibited a strong positive correlation with precision (*ρ* = 0.601, *p* = 3.11 *×* 10^−7^), indicating that targets containing a larger fraction of epitope residues generally yielded higher precision. In contrast, recall showed a moderate negative correlation with epitope density (*ρ* = −0.338, *p* = 0.00764). F1-score showed a weaker but significant positive correlation (*ρ* = 0.310, *p* = 0.0152), whereas MCC showed no significant association with epitope density (*ρ* = −0.031, *p* = 0.812). These results suggest that epitope density affects individual performance metrics differently, with higher epitope density associated with increased precision but reduced recall, rather than producing a uniform improvement across all performance measures.

### S3.2 Impact of Different Features and Modularity on Performance

The CARTnet uses a variety of input features and modified network architecture components. To understand the contribution of each component, we performed an ablation study by retraining the network after removing or replacing individual components. For each ablation setting, the best model was selected based on the AUPRC on the validation dataset, and its performance was subsequently evaluated on the internal test dataset. The results are presented in **Fig. S3** and **Table S5**. The evaluated components include ESM3 structural embeddings, a combination of features such as physicochemical features, secondary structures and sequence collectively referred here as other features, intra-chain contacts derived from the antigen tertiary structure, replacement of the specialized CARTnet block with a standard attention block, and the number of CARTnet blocks.

The results show that the ESM3 structural embeddings have the largest impact on the performance of CARTnet. Removing the ESM3 embeddings resulted in a 31.11% reduction in F1 score, substantially larger than the effects of the other ablations. Removing the intra-chain contacts reduced the F1 score by 5.07%, while replacing the specialized CARTnet block with a standard attention block resulted in a 2.86% reduction. Removing the other features decreased the F1 score by 2.53%, whereas reducing the architecture to a single CARTnet block resulted in the smallest reduction of 1.58%.

A broadly similar trend was observed across the other evaluation metrics. AUPRC, however, remained relatively stable across most ablation settings, with a substantial decrease observed primarily when the ESM3 structural embeddings were removed. Overall, these results indicate that all evaluated components contribute to the predictive performance of CARTnet to varying degrees, with the ESM3 structural embeddings providing the largest contribution and the number of CARTnet blocks having the smallest observed effect.

**Table S5:** Ablation study of the proposed model.

| **Model** | **Acc.** | **Prec.** | **Recall** | **F1** | **AUC** | **AUPRC** | **Prec@Top-10** | **MCC** |
|---|---|---|---|---|---|---|---|---|
| No intrachain | 0.747 | 0.284 | 0.392 | 0.299 | **0.706** | 0.246 | 0.343 | 0.174 |
| No ***F***<sub>res</sub> | 0.750 | 0.292 | 0.401 | 0.307 | 0.702 | 0.250 | 0.328 | 0.185 |
| No ***F***<sub>ESM</sub> | 0.711 | 0.206 | 0.278 | 0.217 | 0.592 | 0.156 | 0.202 | 0.061 |
| Vanilla attention | 0.749 | 0.290 | 0.405 | 0.306 | 0.696 | 0.240 | 0.331 | 0.184 |
| 1 block | 0.752 | 0.297 | 0.401 | 0.310 | 0.704 | **0.251** | 0.330 | 0.189 |
| **Full model** | **0.754** | **0.301** | **0.411** | **0.315** | 0.700 | 0.243 | **0.344** | **0.196** |

**Fig. S3:**
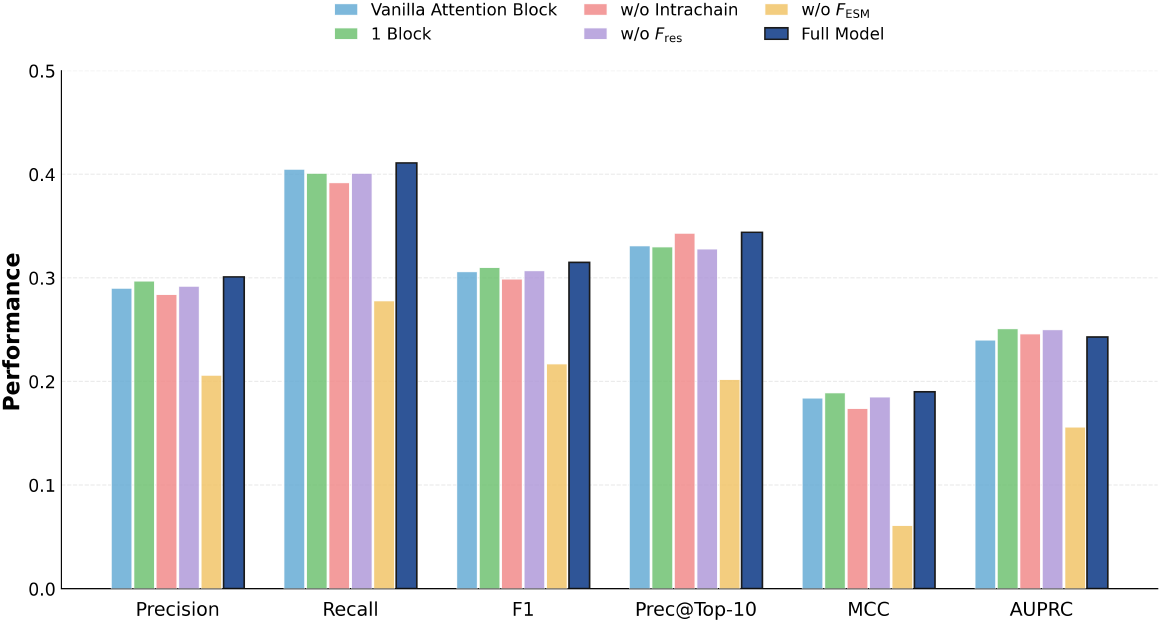
Impact of different groups of features and components on the precision, recall, F1, MCC of the top-20% prediction, precision @ Top-10, and global AUPRC of the test dataset

### S3.3 Impact and Robustness of Template-Guided Prediction

**Table S6:** Displaying the impact of increasing template information on the performance of CARTnet.

| **Temp** | **Acc.** | **Prec.** | **Recall** | **F1** | **AUC** | **AUPRC** | **Top-10** | **MCC** |
|---|---|---|---|---|---|---|---|---|
| 0 | 0.749 | 0.289 | 0.396 | 0.303 | 0.677 | 0.225 | 0.352 | 0.181 |
| 1 | 0.801 | 0.408 | 0.598 | 0.442 | 0.784 | 0.403 | 0.611 | 0.364 |
| 2 | 0.821 | 0.447 | 0.678 | 0.491 | 0.829 | 0.477 | 0.690 | 0.431 |
| 3 | 0.828 | 0.452 | 0.712 | 0.504 | 0.854 | 0.498 | 0.689 | 0.453 |
| 4 | 0.833 | 0.451 | 0.732 | 0.508 | 0.870 | 0.508 | 0.693 | 0.463 |
| 5 | 0.838 | 0.451 | 0.755 | 0.512 | 0.874 | 0.512 | 0.707 | 0.474 |
| 6 | 0.839 | 0.445 | 0.768 | 0.510 | 0.874 | 0.509 | 0.715 | 0.477 |
| 7 | 0.840 | 0.435 | 0.762 | 0.504 | 0.876 | 0.507 | 0.702 | 0.473 |
| 8 | 0.840 | 0.421 | 0.770 | 0.495 | 0.875 | 0.496 | 0.684 | 0.467 |
| 9 | 0.840 | 0.409 | 0.779 | 0.487 | 0.882 | 0.495 | 0.679 | 0.463 |

